# Sucrose overconsumption impairs feeding circuit dynamics and promotes palatable food intake

**DOI:** 10.1101/2023.06.15.545110

**Authors:** Carolyn M. Lorch, Nikolas W. Hayes, Jessica L. Xia, Stefan W. Fleps, Hayley E. McMorrow, Haley S. Province, Joshua A. Frydman, Jones G. Parker, Lisa R. Beutler

## Abstract

Rapid gut-brain communication is critical to maintain energy balance and is disrupted in diet-induced obesity through mechanisms that remain obscure. Specifically, the role of carbohydrate overconsumption in the regulation of interoceptive circuits has been minimally examined *in vivo*. Here we report that an obesogenic high-sucrose diet (HSD) selectively blunts silencing of hunger-promoting AgRP neurons following intragastric delivery of glucose, whereas we previously showed that overconsumption of a high-fat diet (HFD) selectively attenuates lipid-induced neural silencing. By contrast, both HSD and HFD reversibly dampen rapid AgRP neuron sensory inhibition following chow presentation and promote intake of more palatable foods. Our findings reveal that excess sugar and fat pathologically modulate feeding circuit activity in both macronutrient-dependent and -independent ways, and thus may additively exacerbate obesity.

## Introduction

Hunger-promoting, hypothalamic AgRP neurons, located in the arcuate nucleus, monitor energy state and drive appropriate behavioral and physiologic responses to maintain body weight and metabolic health^1–8^. Prior work demonstrates that high-fat diet (HFD)-induced obesity dysregulates AgRP neural responses to nutritional stimuli in a manner that exacerbates palatable food consumption and weight gain^9–12^. However, despite the increasingly clear role of high-sugar foods in driving obesity and metabolic disease, the effects of sugar overconsumption on gut-brain dynamics are largely unknown.

AgRP neurons integrate exterosensory and interoceptive cues to maintain energy homeostasis. AgRP neurons are active in hungry animals, and their stimulation drives feeding and hunger even in conditions where food intake is normally low^3, 4, 13–18^. By contrast, silencing or ablating these neurons suppresses feeding^1, 4, 19^. *In vivo* monitoring of AgRP neuron activity has revealed that their activity is regulated by food-related stimuli across multiple timescales. These neurons are rapidly inhibited when hungry mice are presented with cues that predict imminent food availability^5, 16, 20^. This inhibition precedes consumption, its magnitude correlates with subsequent food intake, and AgRP neuron activity rapidly increases over minutes if animals do not immediately eat^5, 7, 20^. Nutrient sensing in the gastrointestinal (GI) tract is also sufficient to inhibit AgRP neurons proportionally to the caloric content of the intake. This post-ingestive inhibition is mediated by GI distention, and by hormones released from nutrient-sensing enteroendocrine cells that line the proximal small intestine^7–9, 21^. Combined, these signals tune AgRP neuron activity to subserve appropriate feeding behavior and metabolism.

Precisely how diet-induced obesity (DIO) affects the *in vivo* activity of feeding circuits has only recently been explored. Several studies have established that a HFD dysregulates hypothalamic neural responses to nutritional stimuli^9, 10, 22^. In particular, HFD-induced obesity attenuates AgRP neural dynamics in a manner that may counteract weight loss and promote weight regain. Obesity dampens anticipatory chow-mediated AgRP neuron inhibition and reduces intake of chow relative to palatable foods, even prior to substantial weight gain^10^. HFD also attenuates AgRP neuron inhibition in response to nutrient ingestion, an alteration that is selective for dietary fat and lipid-stimulated hormone release^9^. This impairment in nutrient-mediated AgRP neuron inhibition may promote excessive consumption in obese mice, as this inhibition is thought to alleviate the negative affective experience of hunger^20, 23^.

A critical gap in our understanding of how obesogenic diets alter feeding circuit dynamics is whether the changes observed are due to a positive energy balance, availability of palatable food, or dietary macronutrient composition. Recent studies evaluating the effects of DIO on *in vivo* neural dynamics have exclusively employed HFD, making these effects impossible to disentangle^9, 10, 22^. However, the fat-selectivity of the impaired AgRP neuron response to ingested nutrients^9^, together with the recent finding that fats and sugars are sensed by distinct gut-brain pathways^7, 24^, support the idea that diet macronutrient composition may mediate gut-brain axis dysregulation in obesity. By contrast, recent work has shown that, similar to HFD, sugar overconsumption increases the intrinsic activity of AgRP neurons and decreases their sensitivity to the adipokine leptin *ex vivo*^11, 12, 25^. Therefore, obesogenic diets of varying macronutrient composition may also have common effects on feeding circuit activity. Given that obesogenic diets are typically high in both fats and sugars, it is imperative to delineate the contribution of each to pathologic changes in these neural dynamics.

To address how excessive sugar intake modulates *in vivo* feeding circuit activity, we developed an obesogenic high-sucrose diet (HSD), monitored its effects on nutrient-mediated AgRP neuron dynamics and feeding behavior, and determined which changes were reversed upon return to a balanced chow diet. We show that HSD alters AgRP neuron dynamics in both similar and distinct manners to HFD^9, 10^. Like mice on a HFD, HSD-fed mice exhibit reduced chow intake and an associated reduction in AgRP neuron responses to external sensory cues of chow availability. In contrast to mice on a HFD, HSD-fed mice maintain intact AgRP neural responses to intragastrically (IG) infused lipids, but reduced responses to glucose. Taken together, these findings reveal a dissociation between the contributions of calorie excess and macronutrient composition to the neural and behavioral dysregulation that underlie the persistent consequences of obesogenic diets.

## Results

### Liquid sucrose availability alters diet macronutrient composition and increases caloric intake, body weight, and adiposity

To understand the impacts of sugar overconsumption on gut-brain dynamics, we developed a DIO model wherein mice were given *ad libitum* access to a palatable 25% sucrose solution in addition to their regular chow and water. Daily carbohydrate intake was dramatically increased in HSD-fed mice, and total daily caloric intake was modestly augmented compared to controls on a normal chow diet (NCD) (**Fig. 1A-C**). HSD-fed mice gained a significant amount of weight and body fat after four weeks on this diet, and lost a fraction of this weight upon return to a standard chow diet for four weeks (**Fig. 1D-G**). The amount of weight gain observed is less than what has been recently reported in similar approaches using HFD^9, 10^, an observation that is consistent with other models of sugar- and fat-induced obesity^25–27^. These effects were more robust in male compared to female mice (**Fig. S1**). In addition, weight gain was positively correlated with starting weight in the HSD group (**Fig. S2A**).

**Figure 1.**
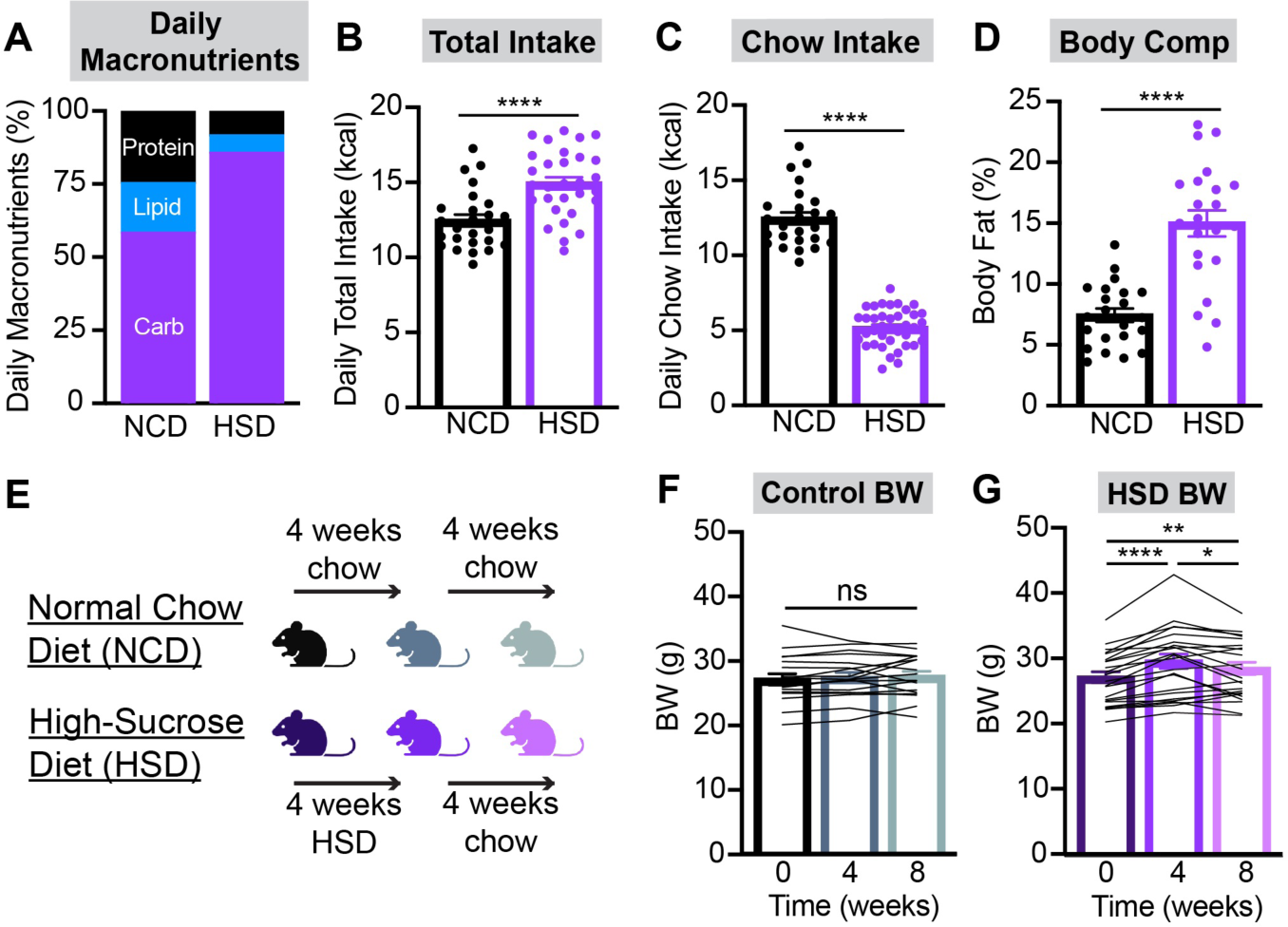
Liquid sucrose availability alters diet macronutrient composition and increases caloric intake and body weight. **(A-C)** Macronutrient composition of intake (A), total daily caloric intake (B), and daily calories from chow (C) in control (NCD) and HSD-fed C57BL6/J mice averaged over three weeks. n = 25-38 mice per group. **(D)** Body fat percentage in NCD and HSD mice after four weeks of chow or HSD, respectively. N = 22 mice per group. **(E)** Timeline of diets administered throughout the remainder of this study. **(F,G)** Body weights (BW) of NCD (F) and HSD (G) mice at baseline, after four weeks of chow or HSD, and after an additional four weeks of chow. n = 18-23 mice per group. *p<0.05, **p<0.01, and ****p<0.0001 as indicated. Baseline body weight was not significantly different between control and HSD mice. (B-D,F,G) Dots or lines represent individual mice. Error bars indicate mean ± SEM.

### High sucrose diet significantly alters hormonal and metabolic parameters

To characterize the physiological consequences of a HSD, we conducted a series of plasma hormone studies and glucose tolerance assays in both NCD and HSD mice at baseline, after 4 weeks of chow or HSD, and after 4 more weeks of chow consumption (**Fig. 1E**). Consistent with a previous report, liquid sucrose overconsumption did not significantly impact fasting plasma insulin levels (**Fig. 2A, B**)^25^, nor did change in insulin levels from baseline to 4 weeks correlate with weight gain (**Fig S2B**). However, our HSD did significantly raise plasma leptin levels compared to control mice (**Figs. 2C-E**). The change in leptin levels from baseline to 4 weeks strongly correlated with weight gain over this time course (**Fig. S2C**).

**Figure 2.**
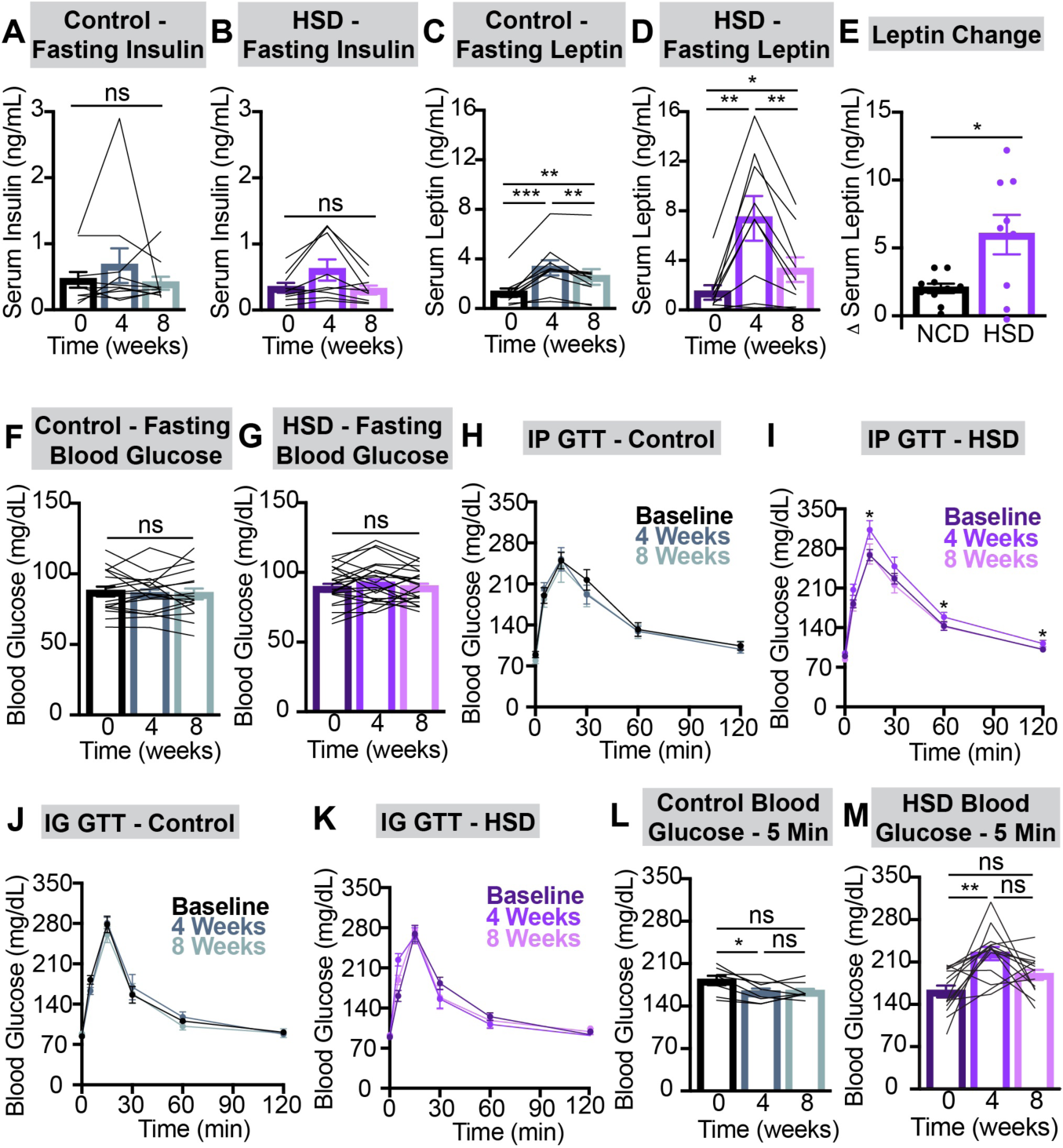
HSD significantly alters glucose homeostasis and circulating leptin. **(A-D)** Plasma insulin (A,B) and leptin (C,D) measured following a six-hour fast in NCD (A,C) and HSD (B,D) mice at baseline, after four weeks of chow or HSD, and after an additional four weeks of chow. n = 9-10 mice per group. **(E)** Change in fasting serum leptin in NCD (C) versus HSD (D) mice from baseline to four weeks. **(F,G)** Blood sugar measured following a six-hour fast in NCD (F) and HSD (G) mice at baseline, after four weeks of chow or HSD, and after an additional four weeks of chow. n = 18-23 mice per group. **(H-K)** Blood glucose following 1.5 mg/g glucose delivered intraperitoneally (H,I) and intragastrically (J,K) in NCD (H,J) and HSD (I,K) mice fasted for six hours at baseline, after four weeks of chow or HSD, and after an additional four weeks of chow. GTT = glucose tolerance test. n = 8-16 mice per group. **(L,M)** Blood glucose values five minutes after intragastric glucose delivery in NCD (L) and HSD (M) mice from (J) and (K), respectively. n = 8-14 mice per group. *p<0.05, **p<0.01, and ***p<0.001 as indicated. (A-G,L,M) Does or lines represent individual mice. Error bars indicate mean ± SEM.

Although fasting blood glucose levels did not significantly change after 4 weeks of NCD or HSD (**Fig. 2F, G**), HSD did impair glucose tolerance. HSD mice demonstrated greater glucose excursions following intraperitoneal (IP) glucose administration after 4 weeks relative to control mice. These changes were rescued upon return to a chow diet (**Fig. 2H, I**). While tolerance to intragastric (IG) glucose was not significantly impaired in HSD mice (**Fig. 2J, K**), they exhibited an early rise in blood glucose levels 5 minutes following glucose delivery not seen in control mice, which was ameliorated upon return to normal chow diet (**Fig. 2L, M**). This early post-ingestive hyperglycemia has been observed previously on a high-sugar diet, and may be due to an altered incretin effect in HSD-fed mice^28^.

The livers of mice after 4 weeks of HSD were steatotic relative to those of control mice (**Fig. S3A-C**). This ectopic fat accumulation was consistent with the elevated fat mass in HSD mice (**Fig. 1D**). However, liver weights (**Fig. S3D**), cholesterol levels (**Fig. S3I**), and plasma liver function indicators including alanine aminotransferase, albumin, alkaline phosphatase, and total bilirubin were not significantly different after 4 weeks of NCD versus HSD (**Fig S3E-H**). Taken together, these results reveal that while 4 weeks of HSD leads to only modest weight gain, sucrose overconsumption causes both glucose intolerance and steatosis, canonical signs of the metabolic syndrome.

### HSD reversibly suppresses fasting-induced hyperphagia and AgRP neuron responses to chow presentation

To investigate the impact of HSD on feeding behavior, we re-fed 6-hour-fasted control and HSD mice with either chow, chocolate, or liquid sucrose (25%). Similar to mice fed a HFD^9, 10^, HSD mice consumed significantly less regular chow relative to baseline fast re-feeding, an effect that was reversed after being replaced on a regular chow diet for 4 weeks (**Fig. 3A, B**). This normalization occurred even though the HSD-fed mice had not completely returned to their baseline body weight (**Fig. 1G**). Chow intake in fast re-fed control mice was consistent across this time course. By contrast, when HSD mice were fasted and re-fed with sucrose, their intake was modestly increased from baseline (**Fig. 3E, F**). Fast re-feeding with chocolate, a palatable food with sugar content intermediate between chow and sucrose solution, was suppressed after 4 weeks of HSD, but to a lesser degree than that of normal chow (**Fig. 3C, D**). Altogether, we found that there was a graded re-feeding response in HSD-fed (but not control) mice that correlated with the sucrose content of the food presented (**Fig. 3G, H**). These results indicate a selective preservation of appetite for sugary food in HSD-fed mice.

**Figure 3.**
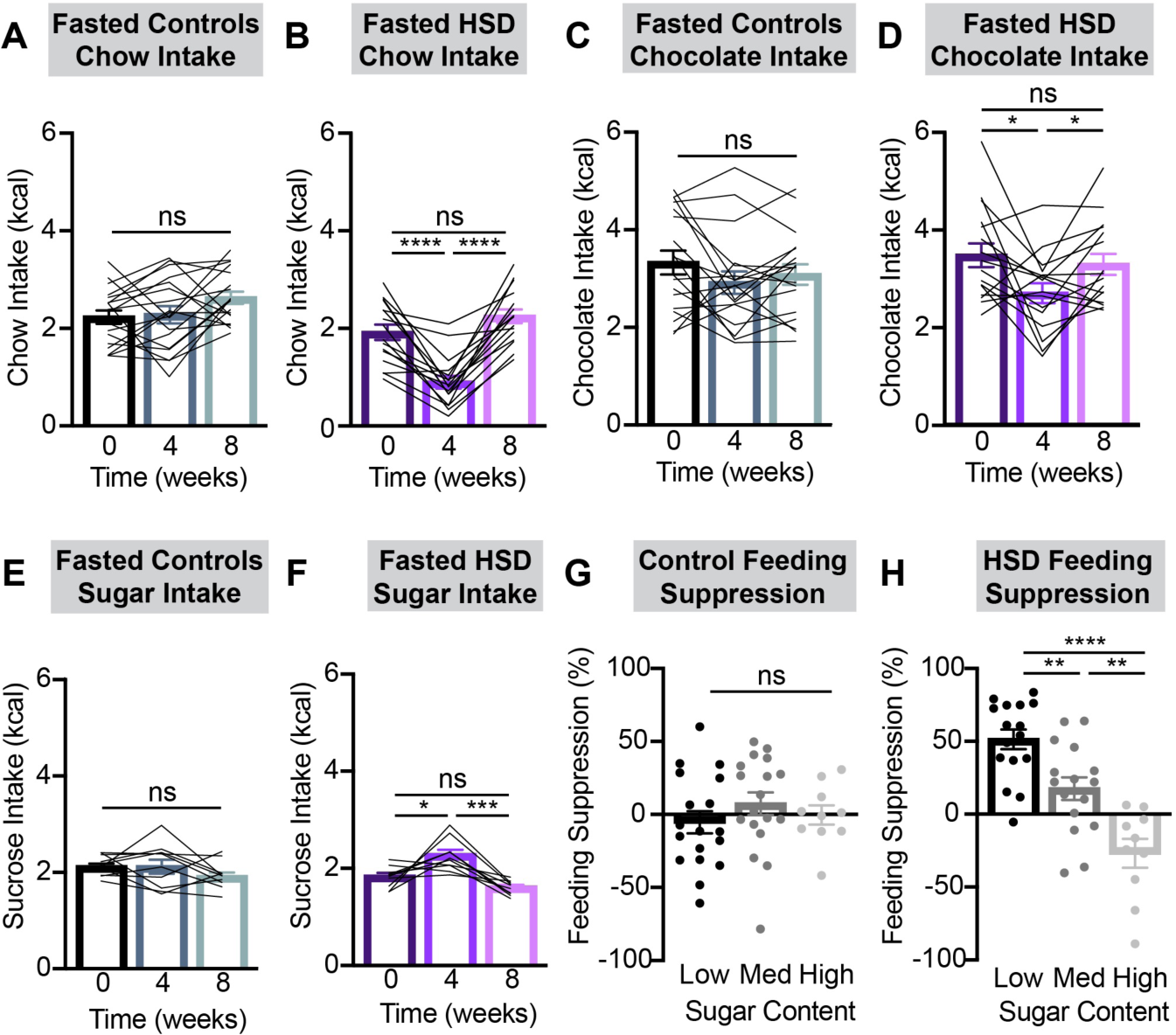
HSD selectively suppresses consumption of low-sugar foods. **(A-F)** Two-hour chow (A,B), chocolate (C,D), or sucrose (E,F) intake following a six-hour fast in NCD (A,C,E) and HSD (B,D,F) animals at baseline, after four weeks of chow or HSD, and after an additional four weeks of chow. n = 10-18 mice per group. **(G,H)** Extent of feeding suppression after four weeks of NCD (G) or HSD (H) relative to baseline. Feeding suppression = ((baseline food intake - four week food intake)/(baseline food intake))*100. Regular chow, chocolate, and 25% sucrose in water represent low-, medium (med)- and high-sugar foods, respectively. n = 10-18 mice per group. *p<0.05, **p<0.01, ***p<0.001, and ****p<0.0001 as indicated. (A-H) Dots or lines represent individual mice. Error bars indicate mean ± SEM.

Given the well-established correlation between anticipatory AgRP neuron inhibition and subsequent consumption^5, 9, 10, 20^, we hypothesized that HSD mice would have blunted pre-consummatory AgRP neuron responses to chow presentation. To test this idea, we used fiber photometry to record Ca^2+^ activity in AgRP neurons during fast re-feeding in control and HSD mice. Indeed, HSD mice had dramatically attenuated AgRP neural inhibition upon chow presentation after 4 weeks of sucrose overconsumption, which is restored following return to a NCD (**Fig. 4B, D-G**). By contrast, mice fed a NCD throughout this time course maintained consistent AgRP neuron responses to chow presentation (**Fig. 4A, C**). In summary, while in a positive energy balance on HSD, mice selectively forgo nutritious, balanced foods (i.e., chow), and this is reversed upon losing weight following return to a normal chow diet. These robust behavioral adaptations are mirrored in the dynamics of AgRP neurons in a similar manner to what is seen in HFD-fed mice^9, 10^.

**Figure 4.**
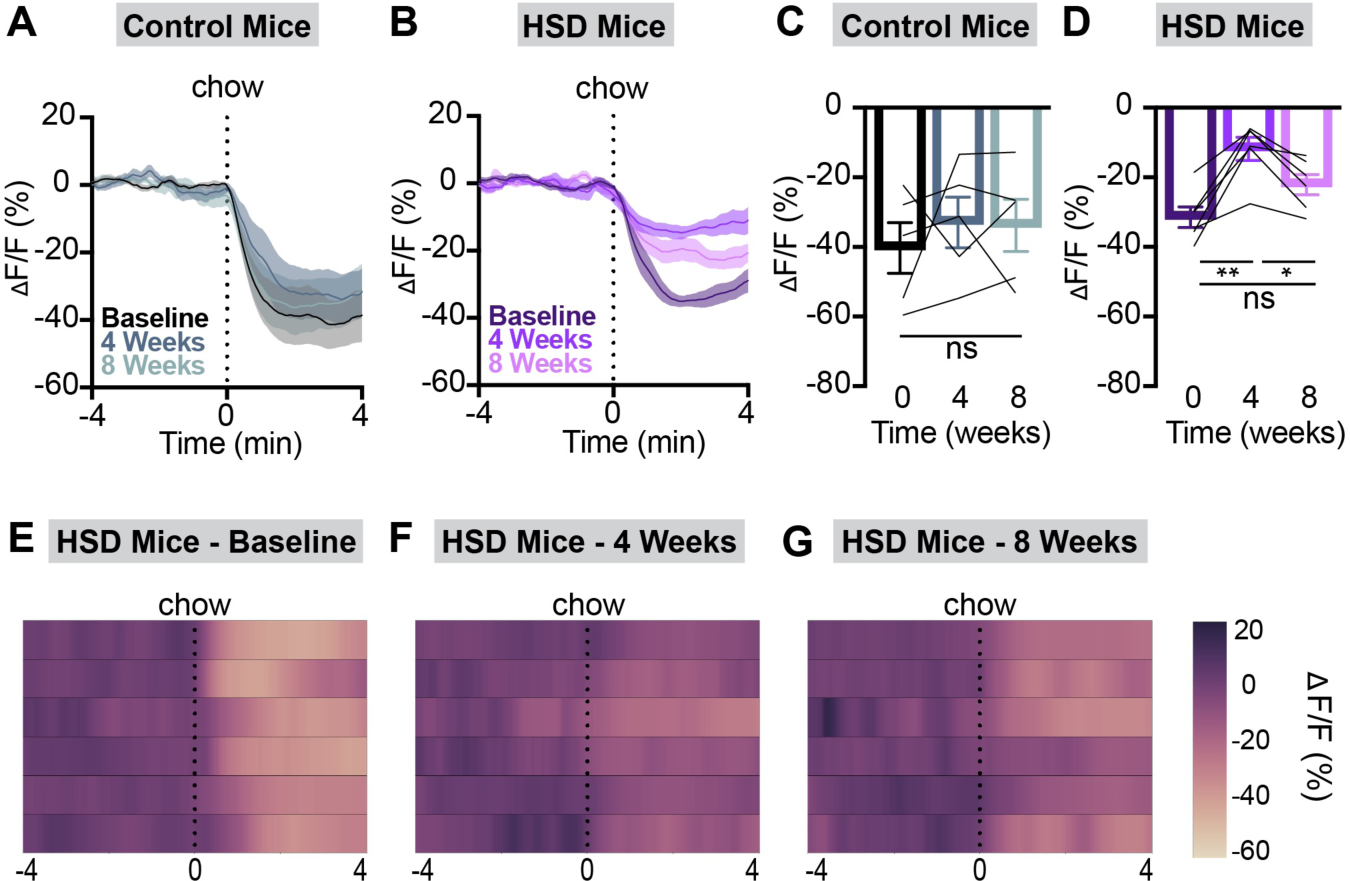
HSD reversibly suppresses AgRP neuron responses to chow. **(A,B)** Calcium signal in AgRP neurons from fasted NCD (A) and HSD (B) mice presented with chow at baseline, after four weeks of chow or HSD (4 weeks), and after an additional four weeks of chow (8 weeks) as indicated. n = 5-6 mice per group. **(C,D)** Average ΔF/F in NCD (C) and HSD (D) mice from (A) and (B) at 2-3 minutes following chow presentation. **(E-G)** Heat map portraying ΔF/F of AgRP neurons in individual HSD mice from (B) and (D) at baseline (E), after four weeks of HSD (F), and after four more weeks of regular chow (G). *p<0.05 and **p<0.01 as indicated. (C,D) Lines represent individual mice. Error bars indicate mean ± SEM.

### High-sucrose diet attenuates AgRP neuron-driven feeding

Because AgRP neural activity is both necessary and sufficient to orchestrate food consumption^1, 3, 4, 15, 16, 19^, we examined whether the ability of AgRP neurons to drive feeding was altered in HSD mice. To do this, we implanted optical fibers above the arcuate nucleus in mice that express channelrhodopsin (ChR2) in AgRP neurons. Our experimental protocol consisted of 30 minutes of habituation followed by 30 minutes of food availability (chow or chocolate) during the presence or absence of light stimulation at baseline and after 4 weeks of HSD (**Fig. 5A**).

**Figure 5.**
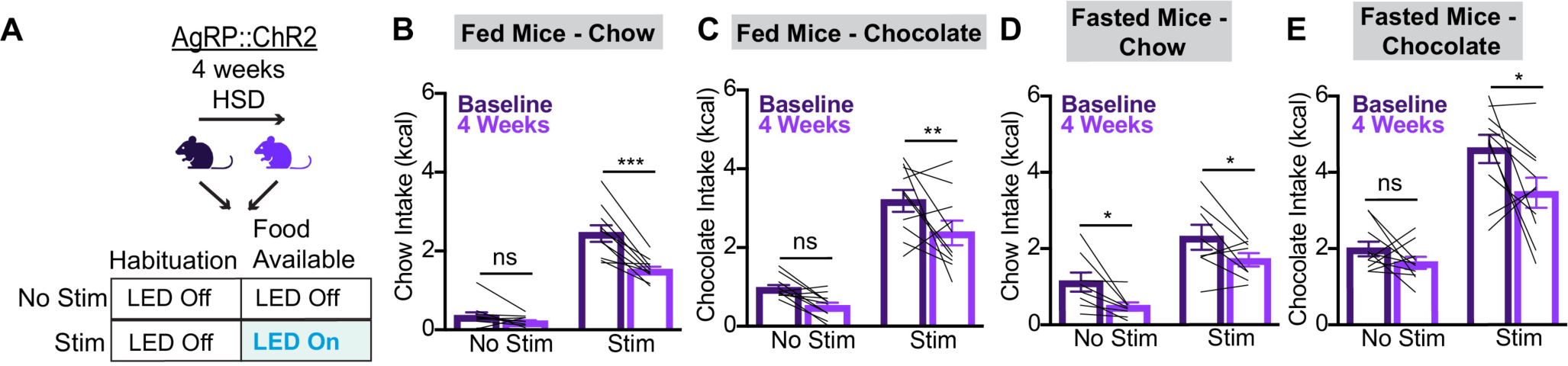
HSD suppresses AgRP neuron-driven food consumption. **(A)** Optogenetic experiment schematic. On separate days, AgRP::ChR2 mice equipped for optogenetic AgRP neuron stimulation were assessed under two protocols. Each session involved a 30-minute habituation period without food and without optical stimulation succeeded by 30 minutes of food availability with (stim) or without (no stim) optical stimulation. Each mouse was tested using both regular chow and chocolate in both the sated and overnight fasted states. These experiments were performed at baseline and repeated after four weeks of *ad libitum* HSD. **(B,C)** Caloric intake of fed mice given chow (B) or chocolate (C) at baseline and after HSD under “stim” and “no stim” protocols. **(D,E)** Caloric intake of fasted mice given chow (D) or chocolate (E) at baseline and after HSD under “stim” and “no stim” protocols. n = 8-10 mice. *p<0.05, **p<0.01, and ***p<0.001 as indicated. In each 2-way ANOVA, p<0.001 for the effect of light stimulation on food intake. (B-E) Lines represent individual mice. Error bars indicate mean ± SEM.

As expected, optogenetic activation of AgRP neurons at baseline strongly increased consumption of both chow and chocolate in sated animals (**Fig. 5B, C**), and modestly increased intake of both chow and chocolate following an overnight fast (**Fig. 5D, E**). After 4 weeks of HSD, the effect of light stimulation on both *ad libitum* feeding and fast re-feeding was attenuated (**Fig. 5B-E**). Control mice lacking ChR2 expression exhibited equivalent feeding behavior with or without light stimulation (**Fig. S4**). Thus, while supraphysiologic AgRP neuron stimulation increases food intake in HSD mice, the effect is less than what is seen in lean mice. Taken together with our fiber photometry results, this finding suggests that, like a HFD, HSD alters both AgRP neuron dynamics and the responsiveness of downstream circuit nodes to AgRP neuron activation ^9, 10^.

### High-sucrose diet selectively and persistently suppresses AgRP neuron responses to glucose

In addition to food presentation, gastric infusion of individual macronutrients is sufficient to durably inhibit AgRP neurons^7, 8^. However, the effects of obesogenic diets on these gut-brain dynamics remain incompletely understood. Given that the AgRP neurons of HFD-induced obese animals become selectively desensitized to dietary fat^9^, we tested the hypothesis that HSD would selectively alter AgRP neuron responses to infused sugars.

To accomplish this, we recorded AgRP neuron activity in HSD and NCD mice in response to a panel of nutrients. AgRP-Cre mice were equipped for fiber photometry and IG nutrient administration via cranial fiberoptic implantation followed by gastric catheterization. HSD and NCD mice were first subjected to baseline recordings wherein isocaloric (1 kcal) and isovolemic (1 mL) solutions of glucose, fructose, sucrose, intralipid, and peptides were infused over a 12-minute period. Water was also administered as a negative control. Intragastric infusion of all nutrients, but not water, reliably inhibited AgRP neurons, consistent with previous reports (**Figs. 6 & S5**)^7–9^. We repeated these nutrient infusions after mice had consumed HSD or NCD for 4 weeks, and a final time after all animals were returned to NCD for 4 weeks. Remarkably, the AgRP neural response to IG glucose in HSD-fed animals was selectively suppressed compared to baseline (**Fig. 6A-G**). Unlike the anticipatory response to chow presentation, the attenuated response to glucose persisted after return to a chow diet and weight loss (**Fig. 6B, D-G**). Responses to fructose, sucrose, lipid, and peptide remained intact in HSD-fed mice, and NCD animals maintained consistent responses to all IG nutrients throughout this time course (**Figs. 6 & S5A-N**). When considered in combination with previously published results from HFD mice^9^, our findings reveal marked differences in the response of interoceptive circuits to the overconsumption of lipid versus carbohydrate-rich obesogenic diets.

**Figure 6.**
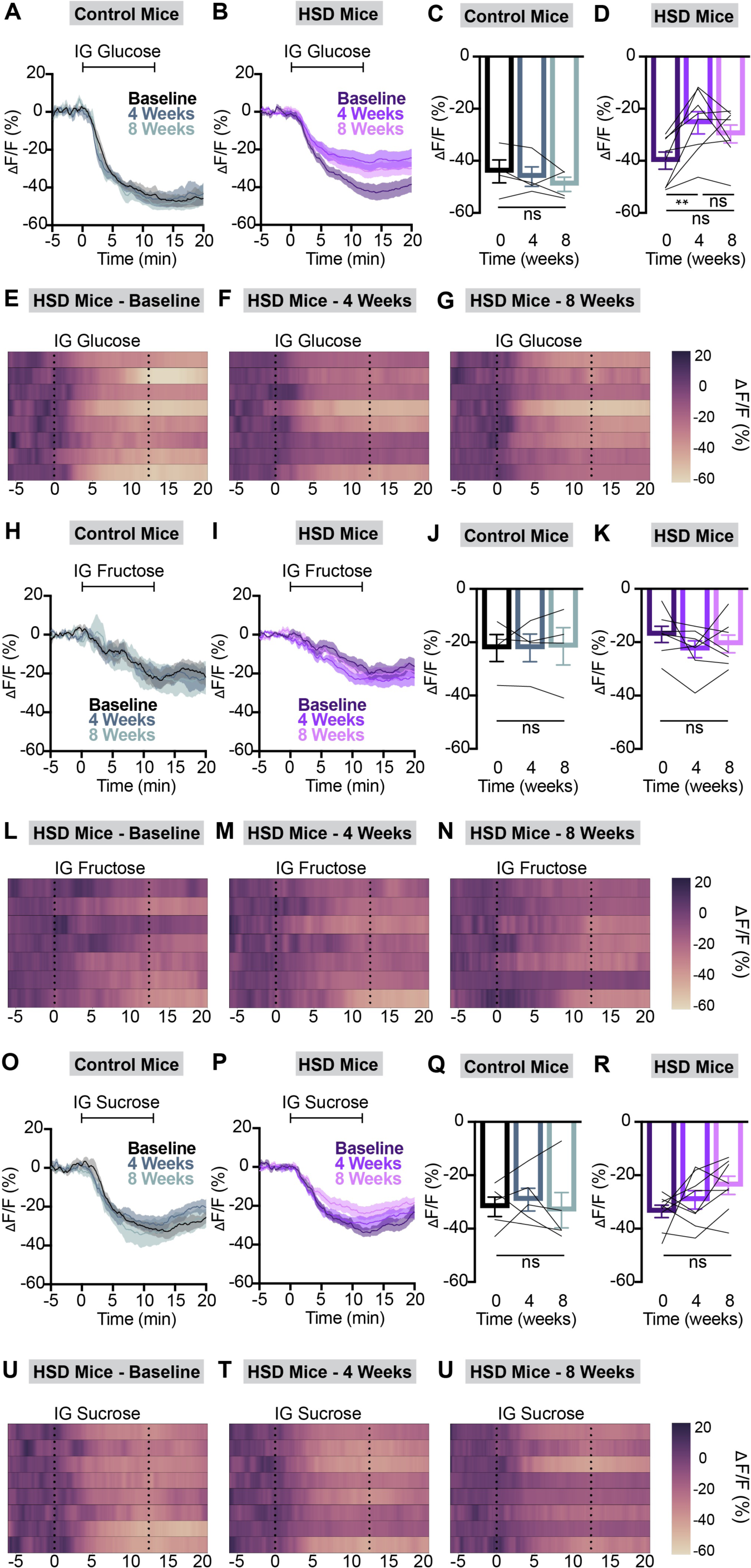
HSD selectively and persistently suppresses AgRP neuron responses to glucose. **(A,B)** Calcium signal in AgRP neurons from fasted NCD (A) and HSD (B) mice during intragastric glucose delivery at baseline, after four weeks of chow or HSD (4 weeks), and after an additional four weeks of chow (8 weeks) as indicated. n = 4-8 mice per group. **(C,D)** Average ΔF/F in NCD (C) and HSD (D) mice from (A) and (B) for one minute surrounding the end of the infusion. **(E-G)** Heat map portraying AgRP neuron fluorescence changes in each HSD mouse from (B) and (D) at baseline (E), after four weeks of HSD (F), and after four more weeks of regular chow (G). **(H,I)** Calcium signal in AgRP neurons from fasted NCD (H) and HSD (I) mice during intragastric fructose delivery at baseline, after four weeks of chow or HSD (4 weeks), and after an additional four weeks of chow (8 weeks) as indicated. n = 4-7 mice per group. **(J,K)** Average ΔF/F in NCD (J) and HSD (K) mice from (H) and (I) for one minute surrounding the end of the infusion. **(L-N)** Heat map portraying AgRP neuron fluorescence changes in each HSD mouse from (I) and (K) at baseline (L), after four weeks of HSD (M), and after four more weeks of regular chow (N). **(O,P)** Calcium signal in AgRP neurons from fasted NCD (O) and HSD (P) mice during intragastric sucrose delivery at baseline, after four weeks of chow or HSD (4 weeks), and after an additional four weeks of chow (8 weeks) as indicated. n = 5-8 mice per group. **(Q,R)** Average ΔF/F in NCD (Q) and HSD (R) mice from (O) and (P) for one minute surrounding the end of the infusion. **(S-U)** Heat map portraying AgRP neuron fluorescence changes in each HSD mouse from (P) and (R) at baseline (S), after four weeks of HSD (T), and after four more weeks of regular chow (U). **p<0.01 as indicated. (C,D,J,K,Q,R) Lines represent individual mice. Error bars indicate mean ± SEM.

## Discussion

Recent studies have begun to unravel how the *in vivo* dynamics of hypothalamic circuits are altered during the development of obesity, producing knowledge that may facilitate novel therapeutic approaches for metabolic diseases. Yet, it is unclear how diet composition impacts the neurophysiological consequences of DIO. Here we leveraged calcium imaging, optogenetics, and a novel rodent model of sucrose-induced obesity to elucidate how dietary macronutrient composition contributes to obesity-induced feeding circuit modulation. Within the context of recently published studies, our results suggest that palatable obesogenic diets lead to macronutrient-dependent and-independent changes in AgRP neuron activity. While altered responses to interoceptive signals depend upon the macronutrient composition of the obesogenic diet, both high-fat and high-sugar diets similarly attenuate the response of AgRP neurons to external sensory cues of food availability. Combined, these changes represent a neural correlate of an altered set point wherein homeostatic feeding circuits are altered by obesity in a manner that defends a new, higher body weight^29, 30^.

### Sucrose overconsumption selectively attenuates glucose-mediated AgRP neuron inhibition and promotes excessive sugar intake

We have previously shown that high-fat DIO leads to selective attenuation of lipid-mediated AgRP neuron inhibition without altering the response of these neurons to ingested glucose or peptides, leading us to interrogate how dietary macronutrient composition modulates obesity-induced changes in gut-brain axis dynamics^9^. Here, we made the parallel observation that a HSD selectively attenuated glucose-induced AgRP neuron inhibition without changing responses to lipids or peptides. As reported previously in HFD-fed mice, this effect persisted even after partial weight loss upon return to a chow diet (**Figs. 6 & S5**). This persistent deficit in glucose-mediated AgRP neural inhibition may compromise the ability of ingested sugar to alleviate the negative valence associated with high AgRP neuron activity, and promote sugar overconsumption^20, 23^. Consistent with this, HSD mice exhibited a stable-to-increased sugar appetite, while their intake of chow and lower-sucrose palatable food appropriately decreased in the obese state (**Fig. 3**). This is analogous to HFD-induced obese mice maintaining a robust appetite for HFD while devaluing chow^10^.

Intriguingly, sucrose overconsumption did not diminish AgRP neuron responses to fructose, the other monosaccharide component of sucrose (**Fig. 6I, K-N**), in HSD-fed mice. Thus, while the response to glucose was dramatically attenuated (**Fig. 6B, D-G**), the stable response to fructose could explain similarly unchanged sucrose-mediated AgRP neuron inhibition (**Fig. 6P, R-U**). Future studies will disentangle how overconsumption of specific monosaccharides contributes to altered nutrient-mediated gut-brain signaling in obesity through monitoring behavioral and neural responses following overconsumption of individual monosaccharides. Additional work will also be required to determine how longer-term hormonal modulators of AgRP neuron activity, feeding, and glucose homeostasis contribute to dysfunctional gut-brain axis dynamics in obesity^31, 32^.

Overall, our findings suggest that diet macronutrient composition plays a more prominent role than caloric intake and body weight in the regulation of gut-brain nutrient-sensing pathways. The persistent desensitization of the gut-brain axis to lipid and glucose following HFD or HSD (**Fig. 6B, D-G**)^9^, respectively, even after weight loss, could contribute to recidivism in obesity, and adds to the body of evidence that dietary history is a critical determinant of future health^29, 30, 33, 34^. Moreover, since typical obesogenic diets are high in both fats and sugars, this work combined with prior HFD findings have laid a foundation for future studies investigating the potentially additive effects of excessive fat and sugar intake on hypothalamic neural dynamics^35–37^.

### Obesity causes macronutrient-independent attenuation of both chow intake and AgRP neuron responses to chow presentation

HFD-induced obesity congruently blunts AgRP neural responses to chow presentation and chow consumption following a fast^9, 10, 38^. These changes are rapidly reversed upon weight loss^9^. Attenuated neural responses to chow and reduced fasting-induced hyperphagia during obesity represent an appropriate adaptation to energy surplus. However, the rapid recovery when animals enter a negative energy balance may promote weight regain. Here we showed that HSD-induced obesity similarly alters the exterosensory regulation of AgRP neuron activity in response to chow presentation (**Figs. 3A, B, & 4**), suggesting that diet palatability and overconsumption, not macronutrient composition, drive this phenomenon. Future work dissecting the effects of overconsumption on AgRP neuron inputs that drive their rapid inhibition and control food intake over multiple time scales will illuminate how obesity modulates the anticipatory response of feeding circuits to food cues^39–42^.

### Obesity blunts AgRP neuron stimulation-induced feeding

Previous studies have shown that HFD-induced obesity attenuates hyperphagia in response to AgRP neuron stimulation^9, 10^. Here, we report a similar effect in HSD mice. Supraphysiologic stimulation of AgRP neurons increased food intake both before and after HSD (**Fig. 5**; *no stim vs. stim*), suggesting that reduced fasting-induced hyperphagia in these animals may be partially due to a failure of food deprivation to maximally activate AgRP neurons in obese mice. This is consistent with earlier findings of reduced fasting- and ghrelin-induced activation of AgRP neurons in obese animals^9, 38, 43, 44^. However, AgRP neuron stimulation-induced food intake in mice during HSD was blunted compared to baseline, which implies reduced sensitivity of downstream circuits to AgRP neuron stimulation (**Fig. 5**; *baseline stim vs. HSD stim*). Taken together, attenuated food intake during AgRP neuron stimulation in obese mice indicates that palatable food overconsumption in these animals is not primarily driven by AgRP neuron activity. Consistent with this, mounting evidence suggests that parallel circuits including mesolimbic dopamine neurons, GABAergic lateral hypothalamic neurons, and non-AgRP expressing arcuate neurons drive feeding independent of energy need^10, 45, 46^.

Taken together, this set of experiments has revealed that DIO exerts macronutrient-dependent blunting of nutrient-mediated gut-brain communication and macronutrient-independent modulation of anticipatory AgRP neuron dynamics. The attenuated pre-consummatory AgRP neuronal response to chow presentation and associated reduction in fasting-induced chow intake is a compensatory adaptation to a sustained positive energy balance on an obesogenic diet. However, this feeding suppression is rapidly reversed before mice return to their baseline weight. This rapid restoration of anticipatory AgRP neuron dynamics and food intake, coupled with persistently blunted macronutrient-mediated AgRP neuron inhibition, promotes excessive palatable food intake, particularly during weight loss, and is likely a key driver of weight regain. Moving forward, characterizing the molecular and circuit-level mechanisms of obesity-induced changes in AgRP neuron dynamics will be critical to identify novel nutritional, hormonal, and neural approaches to permanently treat obesity.

### Limitations of Study

This study has several limitations. First, our HSD is not protein- or vitamin-matched to standard rodent chow, presenting a possible confound when interpreting behavioral and physiological differences between HSD and NCD mice. Future work could employ custom diets which are high in simple sugars but protein- and micronutrient-matched with regular chow to eliminate the possibility that observed changes are due to deficiencies in these other nutrients. However, evidence supports that excessive sugar intake in humans is linked with deficiency in other macro- and micronutrients^47, 48^, making our HSD a physiologically relevant DIO model. Second, while fiber photometry is a powerful method to monitor neural activity, it has caveats. It only captures changes in neural activity over time, and thus offers no insight into how HSD modulates tonic firing of AgRP neurons in intact animals. Nevertheless, recent studies have examined this and found striking similarities with changes seen on a HFD^11, 12, 25^. Finally, fiber photometry cannot capture heterogeneity in neural responses and thus it is unclear whether lipid- and glucose-responsive AgRP neurons are the same or distinct populations. Future work using complementary approaches will address these gaps.

## Acknowledgements

We recognize the Northwestern University Mouse Histology and Phenotyping Laboratory which is supported by NCI P30-CA060553 awarded to the Robert H Lurie Comprehensive Cancer Center for processing mouse liver specimens for histology. We acknowledge Ms. Jiao-Jing Wang from the Northwestern University Microsurgery and Preclinical Research Core for performing a liver function panel using mouse plasma specimens. We thank Dr. Joseph Bass for providing feedback on the manuscript. L.R.B. acknowledges support from the American Diabetes Association Pathway to Stop Diabetes Award. This work was also supported by NIH grants (P30-DK020595), (K08-DK118188), (R01-DK128477) (L.R.B.), and (T32-HL134633) (C.M.L.).

## Author Contributions

C.M.L. and L.R.B. designed and performed experiments, analyzed data, and helped prepare the manuscript. N.W.H. produced custom Python scripts for data analysis. J.G.P. analyzed data and helped prepare the manuscript. J.L.X., S.W.F., H.E.M., H.S.P, and J.A.F. performed experiments.

## STAR Methods

### Animals

Experimental protocols were approved by the Northwestern University IACUC in accordance with the National Institutes of Health guidelines for the Care and Use of Laboratory Animals (Protocol Numbers: IS00015106, IS00023902, and IS00016880). Mice were housed in a 12/12-hour reverse light/dark cycle and given *ad libitum* food and water access. Animals were fed *ad libitum* chow (Envigo, 7012, Teklad LM-495 Mouse/Rat Sterilizable Diet) with or without liquid sucrose (25% sucrose (A15583.0C, Thermo Fisher Scientific) in water (w/v)) for HSD and NCD conditions, respectively. Mice were fasted for either 6 or 16 hours before experiments, as indicated in the text and figures. During this time, they lacked access to chow or sucrose, though they had *ad libitum* water access. *Agrp^tm1(cre)Lowl^*(AgRP-Cre, #012899, Jackson Labs) animals backcrossed onto a C57BL6/J background were used for fiber photometry. For experiments involving optogenetic stimulation of AgRP neurons, AgRP-Cre mice were crossed with 129S-*Gt(ROSA)26Sor^tm32(CAG-COP4*H124R/EYFP)Hze^*(ROSA26-loxStoplox-ChR2-eYFP, #012569, Jackson Labs), yielding double-transgenic mice (AgRP::ChR2). C57BL/6J mice (wildtype #000664, Jackson Labs) were used to measure HSD consumption, as well as the impact of HSD on feeding behavior, glucose homeostasis, hormone levels, and liver anatomy and function. No statistical methods were used to determine sample sizes. Experiments involved male and female mice 2 to 6 months of age unless otherwise indicated. Where sex is not specified in the figure panel, male and female data are combined. Most animals used in feeding or fiber photometry experiments were individually housed to monitor daily caloric intake on HSD or control diets. However, AgRP::ChR2 mice used in optogenetic experiments were group-housed. All experiments were performed during the dark cycle in a dark environment.

### Liquid Sucrose Overconsumption Model

Daily macronutrient consumption, total caloric consumption, and body weight were measured in individually housed mice given pre-measured amounts of a normal chow diet (NCD/controls) or normal chow plus 25% sucrose (HSD). Mice and chow were weighed daily, and volume of 25% sucrose was monitored each morning at the beginning of dark cycle for at least 3 weeks. 25% sucrose was replenished at least every other day.

All baseline fiber photometry recordings, fast re-feeding experiments, assays to determine glucose homeostasis, and hormone measurements were performed on animals fed a standard chow diet. Following these baseline experiments, a subset of mice were placed on HSD. For photometry experiments, animals were assigned to HSD or control groups to match baseline GCaMP signal between groups as closely as possible. Photometry, re-feeding experiments, glucose homeostasis, and hormone assays were performed in HSD and control animals after 3-5 weeks on their respective diets as indicated in text and figures, after which HSD animals were returned to a standard chow diet for 3-5 weeks and these experiments were repeated once more to assess reversibility of observed changes. HSD and control animals underwent the same series of fiber photometry experiments to account for changes in calcium-based fluorescence over time. Optogenetics experiments were performed at baseline and after 3-5 weeks on HSD in the same cohort of mice.

### Stereotaxic Surgery

A recombinant AAV encoding Cre-dependent GCaMP6s (100842-AAV9, AAV9.CAG.Flex.GCaMP6s, Addgene) was employed for fiber photometry studies. The virus was unilaterally injected into the arcuate nucleus (ARC) of AgRP-Cre mice anesthetized with isoflurane. In addition, a photometry cannula (MFC_400/430-0.48_6.3mm_MF2.5_FLT, Doric Lenses) was implanted unilaterally in the ARC at coordinates x = −0.3 mm, y = −1.55 mm, and z = −5.95 mm from bregma. A bronze mating sleeve (SLEEVE_BR_2.5, Doric Lenses) was also adhered to the implant. Mice were given 2-4 weeks for viral expression and surgery recovery before intragastric catheter implantation and/or fiber photometry recordings were commenced.

For optogenetic experiments, we unilaterally inserted fiber optic implants (MFC_200/245_0.37_6.1mm_ZF1.25_FLT, Doric Lenses) above the ARC of AgRP::ChR2 mice anesthetized with isoflurane at coordinates x = −.30, y = −1.55, and z = −5.85 from bregma. Mice were given 1 week to recover prior to experimentation.

Post-operatively, mice were treated with buprenorphine and meloxicam and kept on a heating pad for observation until they were awake and mobile.

### Intragastric Catheter Implantation

Catheterization was performed as previously described^7, 49^. AgRP-Cre mice with correctly placed photometry implants or C57BL6/J mice used for intragastric glucose tolerance testing were anesthetized via ketamine/xylazine. An intrascapular 1-cm skin incision was made, and the skin was dissected from the underlying subcutaneous tissues. Then a 1.5-cm abdominal skin incision was then made caudally from the xyphoid process. The skin was dissected from the underlying peritoneal tissue before an incision was made into the linea alba. A hemostat was used to pierce a small hole through the musculature in the upper back, through which the intragastric catheter was pulled into the abdominal cavity (C30PU-MGA1909, Instech Laboratories). Then the stomach was externalized and punctured, and a purse string suture was used to secure the catheter in the stomach (7-0 USP (Metric 0.5) Nylon, #S-N1718SP13, AD Surgical, Inc.). Absence of leakage was confirmed via saline injection into the catheter. The peritoneum and abdominal skin were closed in two layers (6-0 USP (Metric 0.7) Polypropylene, #S-P618R13, AD Surgical, Inc.). The portion of the catheter extending from between the scapula was secured to a felt button (VABM1B/22, Instech Laboratories) and closed with a magnetic cap (VABM1C, Instech Laboratories). The felt button was sutured to the underlying muscle at the intrascapular site, and the intrascapular skin incision sutured closed (6-0 USP (Metric 0.7) Polypropylene, #S-P618R13, AD Surgical, Inc.). Post-operatively, the mice were treated with enrofloxacin, meloxicam, and buprenorphine and kept on a heating pad for observation until they were awake and mobile. Animals were given a 2-week recovery period before experimentation.

### Fiber Photometry

Two different photometry processors were used for data collection in this study. One setup features LEDs and LED driver separate from the processor (RZ5P, TDT (processor); DC4100 (LED driver); M405FP1 and M470F3 (LEDs), Thorlabs)), while the second setup has these components integrated into the processor (RZ10X, TDT). The neural activity of each mouse was recorded using the same system and patch cord for every session allowing for reliable within-mouse comparisons of calcium signal over time.

Continuous blue LED (465-470 nm) and UV LED (405 nm) served as excitation light sources. These LEDs were modulated at distinct rates and delivered to a filtered minicube (Doric Lenses) before connecting through patch cords (MFP_400/430/1100-0.57_2m_FCM-MF2.5_LAF, Doric Lenses) to mouse implants (MFC_400/430-0.48_6.3mm_MF2.5_FLT, Doric Lenses). GCaMP6s calcium signals and UV isosbestic signals were collected through the same fibers back to dichroic ports of a minicube and transmitted to photoreceivers (Newport Visible Femtowatt Photoreceiver for the RZ5P system; integrated Lux photosensors in the RZ10X system). Digital signals sampled at 1.0173 kHz were then demodulated, lock-in amplified, and collected through the processor (RZ5P or RZ10X, TDT). Data was collected using the software Synapse (TDT), exported, and analyzed using custom Python code.

All fiber photometry experiments were performed in singly-housed AgRP-Cre mice at baseline, after 3-5 weeks of control chow diet or HSD, and after another 3-5 weeks of chow diet. During the experiment, mice were placed in operant chambers (ENV-307W-CT, Med Associates) inside a light- and sound-attenuating cubicle (ENV-022MD, Med Associates) with no food or water access unless otherwise indicated. Mice exhibiting a baseline response to chow presentation of less than ΔF/F = 20% at baseline were deemed technical failures and excluded from further experiments or analysis.

Prior to each recording, the mice’s fiber optic implants were cleaned with 70% ethanol and connector cleaning sticks (MCCS-25, Sticklers) to minimize the potential for debris to interfere with the light path and create a confound. When debris was stuck on the sleeve, a syringe needle was used to remove it.

### Intragastric Infusions

For fiber photometry experiments, a syringe pump (70-2000, Harvard Apparatus) was used to administer nutrients through the intragastric catheters, as has been previously described (cite Beutler et al., 2017, 2020). A total infusion volume of 1 mL was delivered at a constant rate over 12 minutes. Animals were fasted overnight (16 hours) prior to all fiber photometry experiments involving intragastric infusion. The morning of the experiment, the mouse’s intragastric catheter was attached to the syringe pump using polyurethane tubing (VAHBPU-T25, Instech Laboratories) and adaptors (PNP3MC/25, Instech Laboratories; LS25, Instech Laboratories). Mice were given 20 minutes to habituate to the behavioral chambers during photometry recording before initiation of nutrient infusion. Photometry recording continued for another 30 minutes after infusion onset. For peristimulus plots, time zero was defined as when the infusion was initiated. Each nutrient was diluted into deionized water and freshly prepared for each experiment. Intralipid (NDC 0338-0519-09, Fresenius Kabi) was diluted by 50%, while unflavored premium collagen peptides (Sports Research), glucose (G8270, Sigma-Aldrich), fructose (F0060, Tokyo Chemical Industry), and sucrose (Thermo Fisher Scientific) were made into a 25% (w/v) solution. All nutrient infusions were calorie-matched (1 kcal).

For intragastric glucose tolerance tests, glucose (G8270, Sigma-Aldrich) was injected manually as a bolus using a sterile 1 mL Luer Lock syringe (14-955-464, Thermo Fisher Scientific) and an adaptor (VABM1B/22, Instech Laboratories). Glucose (G8270, Sigma-Aldrich) was prepared freshly for each experiment as a 20% (w/v) solution in deionized water and administered at a dose of 1.5 mg/g.

### Food Presentation

To minimize neophobia as a confound, control mice were exposed to chocolate and liquid sucrose once prior to fast re-feeding or photometry experiments where chocolate or sucrose were presented.

For fiber photometry experiments, mice were fasted overnight (16 hours), acclimated to the behavioral chamber for at least 20 minutes, and then presented with chow during fiber photometry recording. For peristimulus plots, time zero was defined as the moment the behavioral chamber was opened.

### Optogenetic Feeding Behavior

Optogenetic experiments were performed in the same AgRP::ChR2 mice both at baseline and after 3-5 weeks of HSD. Mice used in these experiments were group-housed and ranged from 2-12 months old. Mice were given a week to recover after implant surgery. During this time, they were habituated to handling and optogenetic patch cord tethering several times.

An LED source and programmable TTL pulse generator (Prizmatix) was used to generate a train of blue light (460 nm, 2 s ON / 3 s OFF, 10 ms pulse width, 20 Hz, 10-15 mW)^9, 50^. Fiber optic patch cables (500um POF N.A. 0.63 L=75cm, Prizmatix) were connected to mouse implants (MFC_200/245-0.37_6.1mm_ZF1.25_FLT, Doric Lenses) via a zirconia mating sleeve (F210-3001, Doric Lenses).

All experiments involve 30 minutes of habituation without LED stimulation or food availability followed by 30 minutes of access to either chocolate or chow with or without light stimulation. Each light stimulation protocol was performed in both fed and fasted (overnight, 16 hours) states in the same mice on different days, and with access to either chocolate or chow on separate days. For both chocolate and chow, a single, pre-weighed piece of food was placed in the behavior chamber at the end of habituation and weighed once again after the 30 minutes of food access.

### Glucose Tolerance Testing

At baseline, after 3-5 weeks of control chow diet or HSD, and after another 3-5 weeks of chow diet, C57BL6/J mice were fasted for six hours before glucose (G8270, Sigma-Aldrich) administration. Glucose was delivered through either intraperitoneal injection of 15% glucose (w/v) in water or intragastric infusion of 20% glucose (w/v) in water. Either way, glucose was dosed at 1.5 mg/g body weight. Blood glucose concentrations were measured from tail blood using an automatic glucose monitor (FreeStyle Lyte). Fasting baseline blood glucose was measured, and blood glucose concentrations were also recorded at 5, 15, 30, 60, and 120 minutes following glucose administration.

### Plasma Hormone Studies

At baseline, after 3-5 weeks of control chow diet or HSD, and after another 3-5 weeks of chow diet, tail blood was collected from six-hour-fasted C57BL6/J mice in EDTA-coated tubes for plasma separation. Plasma leptin (90030, Crystal Chem) and insulin (90080, Crystal Chem) ELISAs were performed per the manufacturer’s instructions.

### Plasma Liver Panel

After 4 weeks of control chow diet or HSD, *ad libitum-*fed C57BL6/J mice were euthanized before blood collection via cardiac puncture into EDTA-coated tubes for plasma separation. Using these plasma samples, a mammalian liver profile (500-0040, Abaxis) was conducted by the Northwestern University Microsurgery and Preclinical Research Core for the quantitative analysis of albumin (ALB), alanine aminotransferase (ALT), alkaline phosphatase (ALP), cholesterol (CHOL), and total bilirubin (TBIL).

### Liver Histology

After 4 weeks of control chow diet or HSD, *ad libitum-*fed C57BL6/J mice were euthanized and livers were removed, and an approximately 100mg piece from the left lobe was fixed overnight in 4% paraformaldehyde (P6148, Sigma-Aldrich). The next day, the liver was placed in 70% ethanol and stored at room temperature until being processed for histology. The tissue was paraffin embedded, sectioned, and hematoxylin- and eosin-stained, and then imaged for analysis. The extent of hepatic steatosis was quantified using the NAFLD scoring method^51^.

### Quantification and Statistical Analysis

#### Photometry Analysis

Custom Python scripts (https://github.com/nikhayes/fibphoflow) were used to analyze photometry data, and statistical analyses and data visualizations were generated with Prism and Python. Briefly, photometry recordings included AgRP neuron GCaMP activity traces and corresponding UV isosbestic traces, which were smoothed with a low-pass filter and downsampled to 1 Hz. The isosbestic trace was subtracted from the paired GCaMP activity trace to adjust for motion artifact and photobleaching-induced decreases in photometry signal. Normalization of AgRP neuron responses to nutritional stimuli relative to baseline activity was performed via the formula: ΔF/F = (F_t_ – F_0_) / F_0_, where F_t_ represents fluorescence at time (t), and F_0_ represents the average fluorescence during the five-minute baseline period preceding time zero. To determine statistical differences, the average ΔF/F was computed within the time frame indicated in the legend for Figures 4, 6, and S5.

#### Behavior Data Analysis

To determine chow, chocolate, and liquid sucrose consumption for fast re-feeding and optogenetics experiments, food items were weighed manually at the indicated time points.

#### Statistical Analysis

Collection and analysis of fiber photometry data were performed similarly to what has been described previously^7, 9^. For photometry traces depicted in Figures 4, 6, and S5, ΔF/F (%) refers the mean ΔF_t_/F_0_ * 100. Bar graphs quantifying AgRP neuron neural responses to chow and to intragastric nutrients in Figures 4, 6, and S5 represent the average ΔF/F (%) over a 1-minute period following food presentation (**Fig. 4**) or at the end of nutrient infusion (**Figs. 6 & S5**). The time frame represented for each study is specified in the figure legends.

The effects of HSD or NCD and eventual return to a balanced diet on absolute body weight were analyzed via one-way, repeated-measures ANOVA (**Figs. 1 & S1**). Total daily caloric intake, daily caloric intake from chow, and body fat mass percentage in HSD vs NCD mice were assessed via unpaired T-tests (**Figs. 1 & S1**). The effects of HSD or NCD and subsequent return to chow diet on fasting serum insulin, fasting serum leptin, and fasting blood glucose were analyzed using one-way, repeated-measures ANOVA (**Fig. 2**). The effects of HSD versus NCD on the change in fasting serum leptin levels (ng/mL) from baseline to 4 weeks were analyzed using an unpaired T-test (**Fig. 2**). The effects of NCD or HSD on IG or IP glucose tolerance were assessed via two-way, repeated-measures ANOVA (**Fig. 2**). A Pearson’s correlation test was used to determine the relationship between Δ body weight (g) from baseline to 4 weeks and (1) baseline body weight, (2) change in fasting serum leptin, and (3) change in fasting serum insulin (**Fig. S2**). Liver function and steatosis data were analyzed via unpaired T-tests (**Fig. S3**). The effects of HSD or NCD and subsequent return to chow diet on fast re-feeding of chow, chocolate, and liquid sucrose were analyzed using one-way, repeated-measures ANOVA (**Fig. 3**). The change in fast re-feeding by food category in NCD or HSD mice was analyzed using one-way ANOVA (**Fig. 3**). The effects of HSD or NCD and subsequent return to chow diet on AgRP neuron calcium dynamics in response to chow and intragastrically administered nutrients were analyzed using one-way, repeated-measures ANOVA (**Figs. 4, 6, & S5**). The effects of AgRP neuron optogenetic stimulation on feeding before and after HSD were compared using two-way, repeated-measures ANOVA (**Figs. 5 & S4**). The Holm-Sidak multiple comparisons test was used in conjunction with ANOVA where appropriate. Prism was used for all statistical analysis. Sample sizes are specified in figure legends. Significance was defined as p<0.05 and is specified on figures and in figure legends. When multiple technical replicates of a given experiment were conducted, these trials were averaged and handled as one biological replicate for the purposes of data analysis and visualization.

---

**Figure S1.**
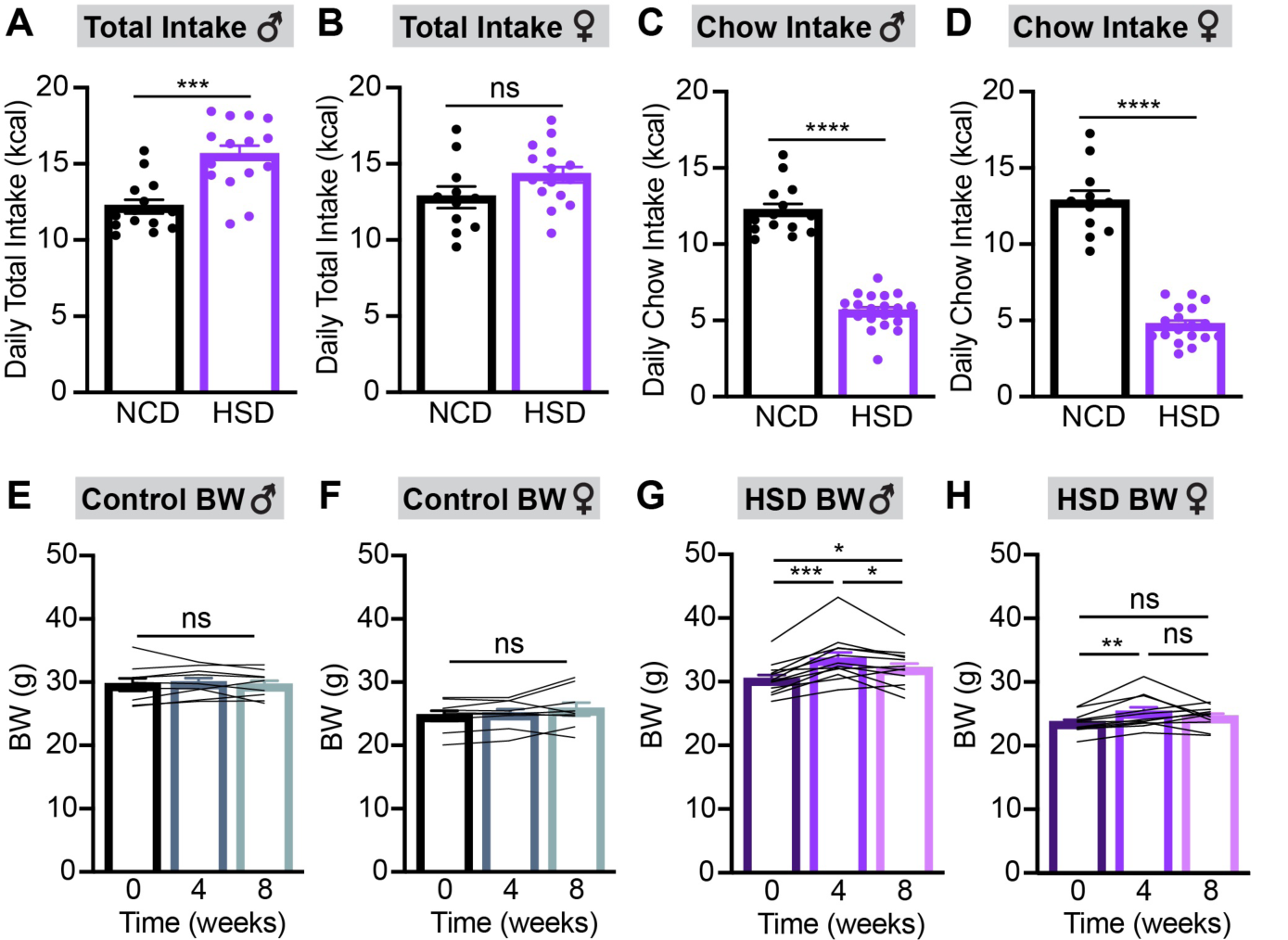
Liquid sucrose availability has sex-specific impacts on caloric intake and body weight. **(A-D)** Total daily caloric intake (A,B) and daily calories from chow (C,D) in control (NCD) and HSD male (A,C) and female (B,D) mice averaged over three weeks. n = 11-20 mice per group. **(E-H)** Body weights in NCD male (E) and female (F) and HSD male (G) and female (H) mice at baseline, after four weeks of chow or HSD, and after an additional four weeks of chow. n = 9-12 mice per group. *p<0.05, **p<0.01, ***p<0.001, and ****p<0.0001 as indicated. Baseline body weight was not significantly different between control and HSD mice of the same sex. (A-H) Dots or lines represent individual mice. Error bars indicate mean ± SEM.

**Figure S2.**
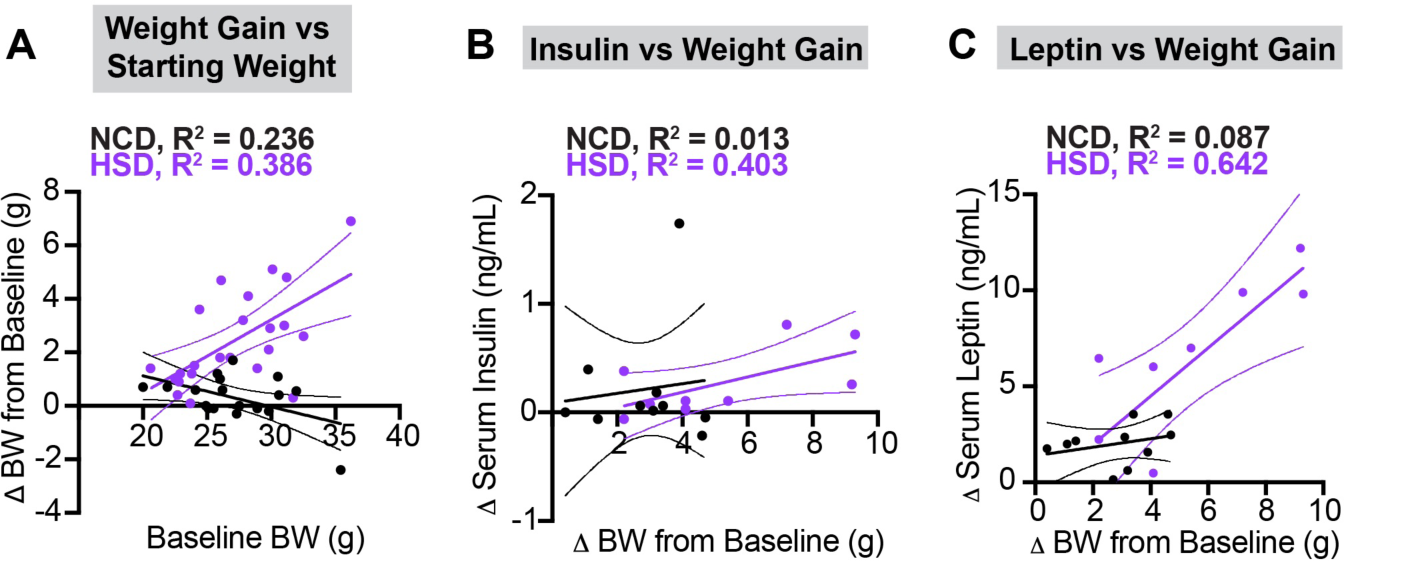
Weight gain on HSD is associated with higher baseline body weight and serum leptin levels. **(A)** Baseline weight versus weight gain in NCD (black, R^2^ = 0.236, p = 0.041) and HSD (purple, R^2^ = 0.386, p=0.002) after four weeks of chow or HSD, respectively. n = 18-23 mice per group. **(B)** Weight gain versus fasting serum insulin change in NCD (black, R^2^ = 0.013, p = 0.750) and HSD (purple, R^2^ = 0.403, p=0.066) after four weeks of chow or HSD, respectively. n = 9-10 mice per group. **(C)** Weight gain versus fasting serum leptin change in NCD (black, R^2^ = 0.087, p = 0.408) and HSD (purple, R^2^ = 0.642, p=0.009) after four weeks of chow or HSD, respectively. n = 9-10 mice per group. Dots represent individual mice. Error bars indicate mean slope ± 95% CI.

**Figure S3.**
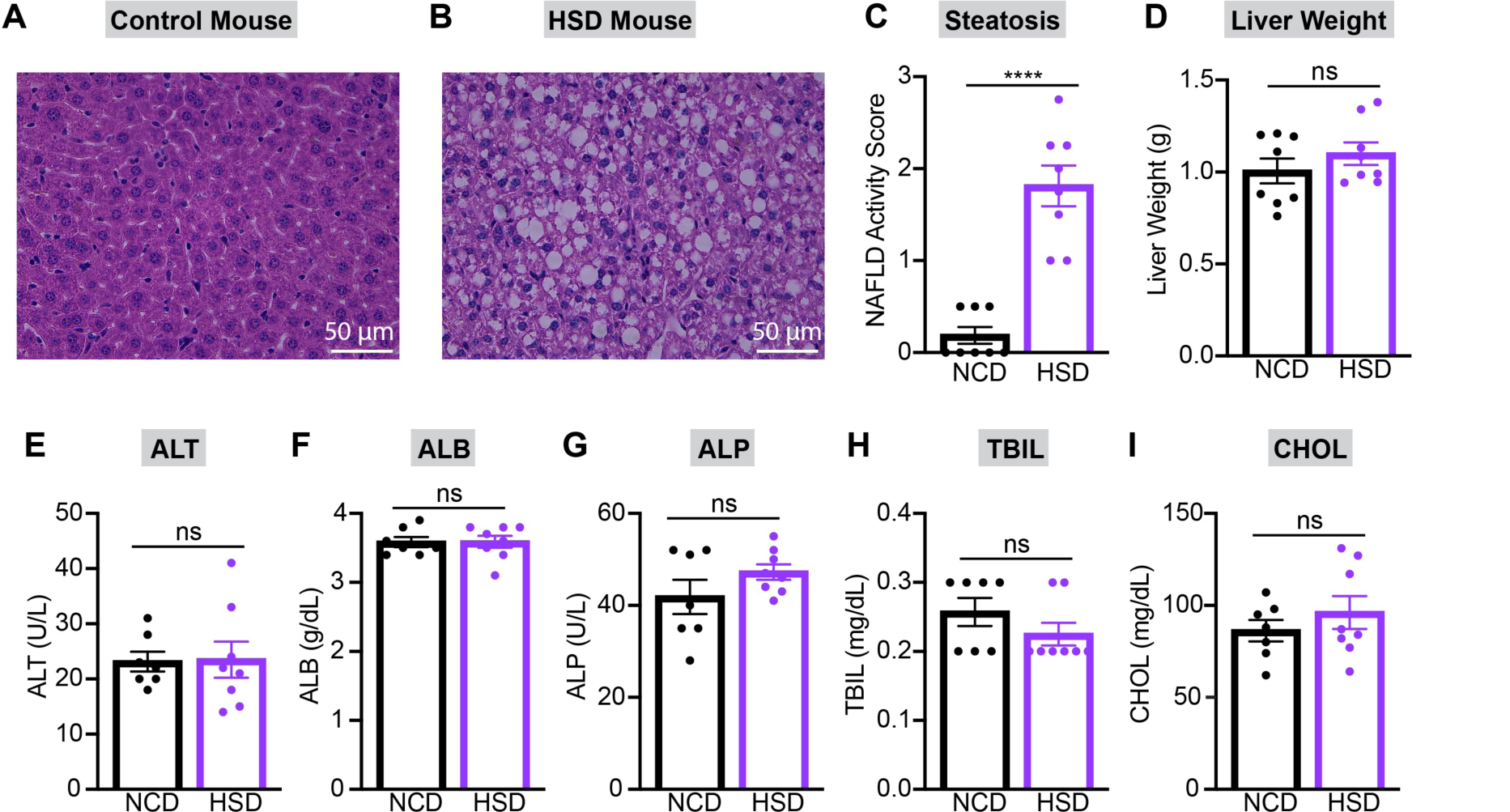
HSD causes hepatic steatosis but does not impair liver function after four weeks. **(A,B)** Representative hematoxylin and eosin-stained liver sections from NCD (A) and HSD (B) mice after four weeks of chow or HSD. **(C)** Percentage of images consisting of lipid droplets in liver sections from NCD versus HSD mice after 4 weeks of chow or HSD, respectively. n = 8 mice per group. **(D)** Weight of livers in *ad libitum*-fed NCD and HSD mice after four weeks of chow or HSD, respectively. n = 8 mice per group. **(E-I)** Alanine transaminase (ALT) (E), albumin (ALB) (F), alkaline phosphatase (ALP) (G), total bilirubin (TBIL) (H), and cholesterol (CHOL) (I) levels in plasma from *ad libitum*-fed NCD and HSD mice after four weeks of chow or HSD, respectively. n = 7-8 mice per group. ***p<0.001 as indicated. (C-I) Dots represent individual mice. Error bars indicate mean ± SEM.

**Figure S4.**
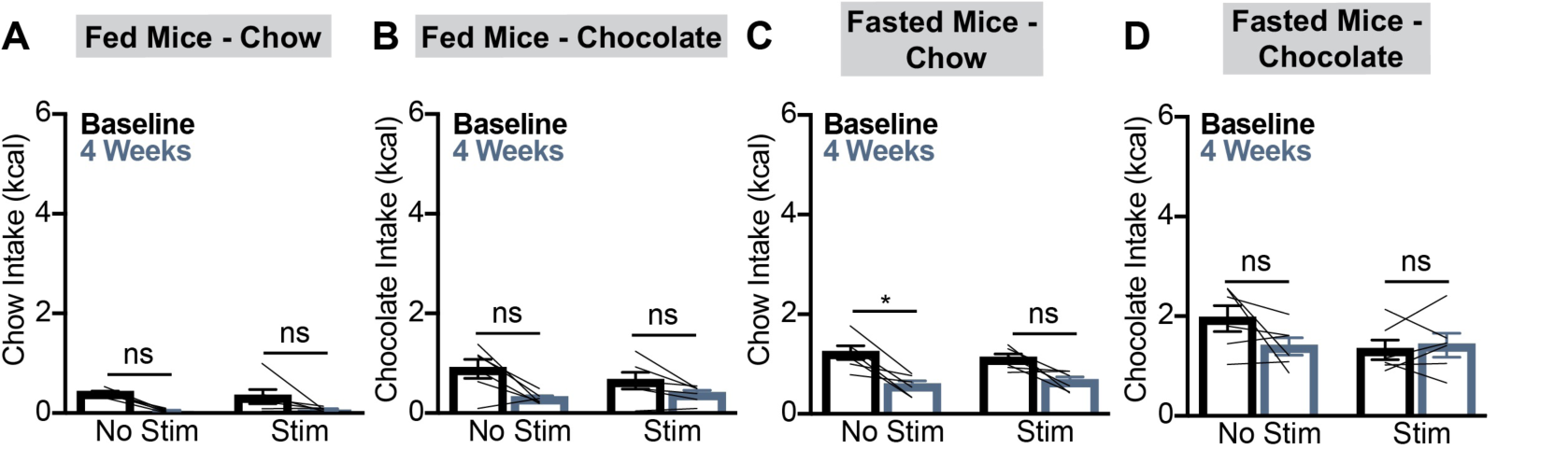
Blue light stimulation does not impact food intake. **(A,B)** Caloric intake of fed mice given chow (A) or chocolate (B) at baseline and after HSD under “stim” and “no stim” protocols. **(C,D)** Caloric intake of fasted mice given chow (C) or chocolate (D) at baseline and after HSD under “stim” and “no stim” protocols. n = 6 mice. *p<0.05 as indicated. In each 2-way ANOVA, p>0.05 for the effect of light stimulation on food intake. (A-D) Lines represent individual mice. Error bars indicate mean ± SEM.

**Figure S5.**
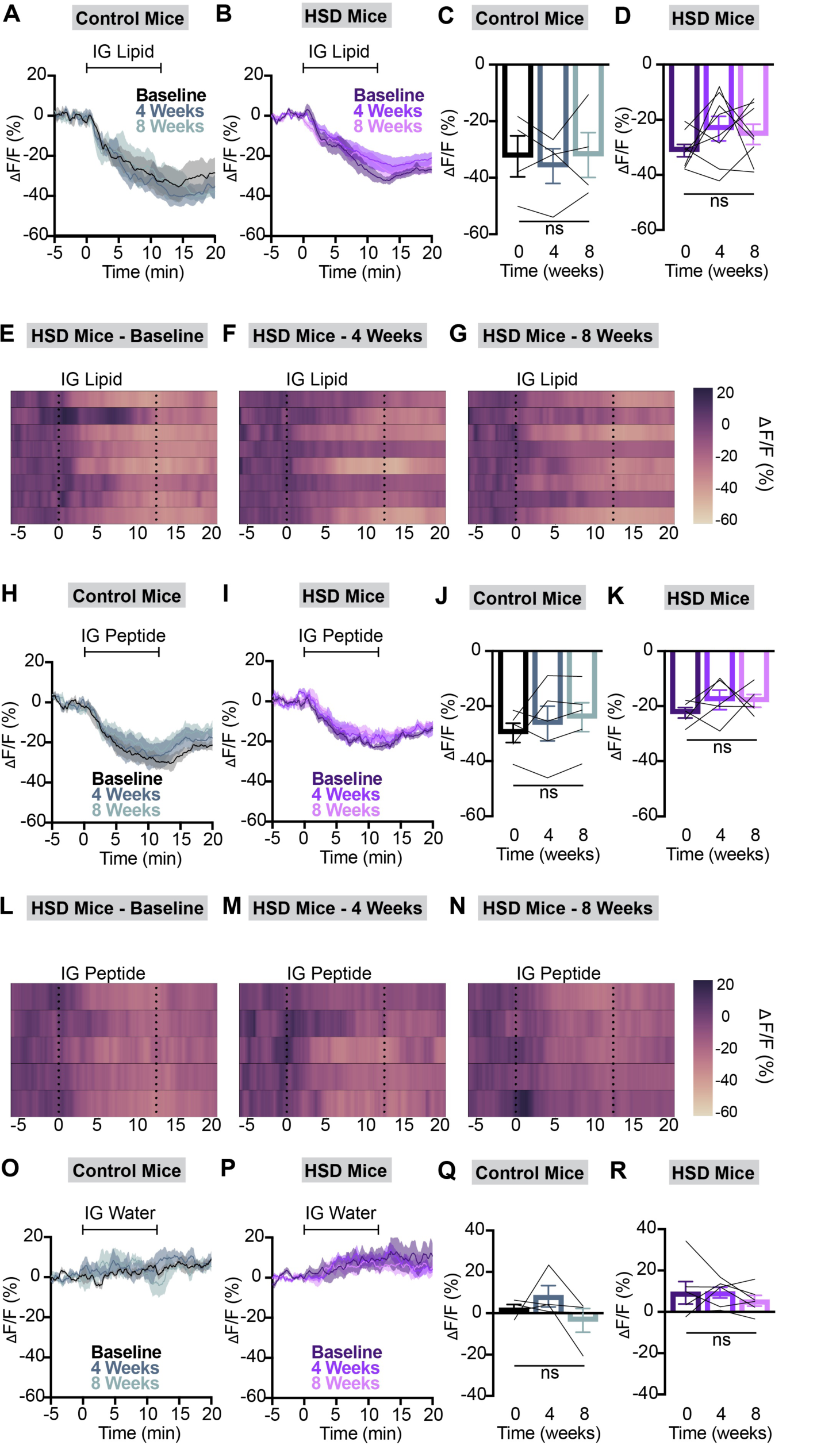
HSD does not alter AgRP neuron responses to lipids or peptides. **(A,B)** Calcium signal in AgRP neurons from fasted NCD (A) and HSD (B) mice during intragastric intralipid delivery at baseline, after four weeks of chow or HSD (4 weeks), and after an additional four weeks of chow (8 weeks) as indicated. n = 4-8 mice per group. **(C,D)** Average ΔF/F in NCD (C) and HSD (D) mice from (A) and (B) for one minute surrounding the end of the infusion. **(E-G)** Heat map portraying AgRP neuron fluorescence changes in each HSD mouse from (B) and (D) at baseline (E), after four weeks of HSD (F), and after four more weeks of regular chow (G). **(H,I)** Calcium signal in AgRP neurons from fasted NCD (H) and HSD (I) mice during intragastric peptide delivery at baseline, after four weeks of chow or HSD (4 weeks), and after an additional four weeks of chow (8 weeks) as indicated. n = 5 mice per group. **(J,K)** Average ΔF/F in NCD (J) and HSD (K) mice from (H) and (I) for one minute surrounding the end of the infusion. **(L-N)** Heat map portraying AgRP neuron fluorescence changes in each HSD mouse from (I) and (K) at baseline (L), after four weeks of HSD (M), and after four more weeks of regular chow (N). **(O,P)** Calcium signal in AgRP neurons from fasted NCD (O) and HSD (P) mice during intragastric water delivery at baseline, after four weeks of chow or HSD (4 weeks), and after an additional four weeks of chow (8 weeks) as indicated. n = 4-6 mice per group. **(Q,R)** Average ΔF/F in NCD (Q) and HSD (R) mice from (O) and (P) for one minute surrounding the end of the infusion. (C,D,J,K,Q,R) Lines represent individual mice. Error bars indicate mean ± SEM.

## References

1. Luquet, S., Perez, F.A., Hnasko, T.S., and Palmiter, R.D. (2005). NPY/AgRP Neurons Are Essential for Feeding in Adult Mice but Can Be Ablated in Neonates. Science 310, 683–685. doi:10.1126/science.1115524.

2. Konner, A.C., Janoschek, R., Plum, L., Jordan, S.D., Rother, E., Ma, X., Xu, C., Enriori, P., Hampel, B., Barsh, G.S., et al. (2007). Insulin action in AgRP-expressing neurons is required for suppression of hepatic glucose production. Cell metabolism 5, 438–449. S1550-4131(07)00131-3 [pii].

3. Aponte, Y., Atasoy, D., and Sternson, S.M. (2011). AGRP neurons are sufficient to orchestrate feeding behavior rapidly and without training. Nature neuroscience 14, 351–355. 10.1038/nn.2739 [doi].

4. Krashes, M.J., Koda, S., Ye, C., Rogan, S.C., Adams, A.C., Cusher, D.S., Maratos-Flier, E., Roth, B.L., and Lowell, B.B. (2011). Rapid, reversible activation of AgRP neurons drives feeding behavior in mice. The Journal of clinical investigation 121, 1424–1428. 10.1172/JCI46229 [doi].

5. Chen, Y., Lin, Y.C., Kuo, T.W., and Knight, Z.A. (2015). Sensory detection of food rapidly modulates arcuate feeding circuits. Cell 160, 829–841. 10.1016/j.cell.2015.01.033 [doi].

6. Steculorum, S.M., Ruud, J., Karakasilioti, I., Backes, H., Engstrom Ruud, L., Timper, K., Hess, M.E., Tsaousidou, E., Mauer, J., Vogt, M.C., et al. (2016). AgRP Neurons Control Systemic Insulin Sensitivity via Myostatin Expression in Brown Adipose Tissue. Cell 165, 125–138. S0092-8674(16)30194-5 [pii].

7. Beutler, L.R., Chen, Y., Ahn, J.S., Lin, Y.-C., Essner, R.A., and Knight, Z.A. (2017). Dynamics of Gut-Brain Communication Underlying Hunger. Neuron 96, 461–475.e465. https://doi.org/10.1016/j.neuron.2017.09.043.

8. Su, Z., Alhadeff, A.L., and Betley, J.N. (2017). Nutritive, Post-ingestive Signals Are the Primary Regulators of AgRP Neuron Activity. Cell reports 21, 2724–2736. S2211-1247(17)31673-X [pii].

9. Beutler, L.R., Corpuz, T.V., Ahn, J.S., Kosar, S., Song, W., Chen, Y., and Knight, Z.A. (2020). Obesity causes selective and long-lasting desensitization of AgRP neurons to dietary fat. eLife 9, 10.7554/eLife.55909. e55909 [pii].

10. Mazzone, C.M., Liang-Guallpa, J., Li, C., Wolcott, N.S., Boone, M.H., Southern, M., Kobzar, N.P., Salgado, I.A., Reddy, D.M., Sun, F., et al. (2020). High-fat food biases hypothalamic and mesolimbic expression of consummatory drives. Nature neuroscience. 10.1038/s41593-020-0684-9 [doi].

11. Baver, S.B., Hope, K., Guyot, S., Bjorbaek, C., Kaczorowski, C., and O’Connell, K.M. (2014). Leptin modulates the intrinsic excitability of AgRP/NPY neurons in the arcuate nucleus of the hypothalamus. The Journal of neuroscience: the official journal of the Society for Neuroscience 34, 5486–5496. 10.1523/JNEUROSCI.4861-12.2014 [doi].

12. Wei, W., Pham, K., Gammons, J.W., Sutherland, D., Liu, Y., Smith, A., Kaczorowski, C.C., and O’Connell, K.M. (2015). Diet composition, not calorie intake, rapidly alters intrinsic excitability of hypothalamic AgRP/NPY neurons in mice. Scientific reports 5, 16810. 10.1038/srep16810 [doi].

13. Hahn, T.M., Breininger, J.F., Baskin, D.G., and Schwartz, M.W. (1998). Coexpression of Agrp and NPY in fasting-activated hypothalamic neurons. Nature neuroscience 1, 271–272. 10.1038/1082 [doi].

14. van den Top, M., Lee, K., Whyment, A.D., Blanks, A.M., and Spanswick, D. (2004). Orexigen-sensitive NPY/AgRP pacemaker neurons in the hypothalamic arcuate nucleus. Nature neuroscience 7, 493–494. 10.1038/nn1226 [doi].

15. Takahashi, K.A., and Cone, R.D. (2005). Fasting induces a large, leptin-dependent increase in the intrinsic action potential frequency of orexigenic arcuate nucleus neuropeptide Y/Agouti-related protein neurons. Endocrinology 146, 1043–1047. en.2004-1397 [pii].

16. Mandelblat-Cerf, Y., Ramesh, R.N., Burgess, C.R., Patella, P., Yang, Z., Lowell, B.B., and Andermann, M.L. (2015). Arcuate hypothalamic AgRP and putative POMC neurons show opposite changes in spiking across multiple timescales. eLife 4, 10.7554/eLife.07122.[doi].

17. Livneh, Y., Ramesh, R.N., Burgess, C.R., Levandowski, K.M., Madara, J.C., Fenselau, H., Goldey, G.J., Diaz, V.E., Jikomes, N., Resch, J.M., et al. (2017). Homeostatic circuits selectively gate food cue responses in insular cortex. Nature 546, 611–616. 10.1038/nature22375 [doi].

18. Essner, R.A., Smith, A.G., Jamnik, A.A., Ryba, A.R., Trutner, Z.D., and Carter, M.E. (2017). AgRP Neurons Can Increase Food Intake during Conditions of Appetite Suppression and Inhibit Anorexigenic Parabrachial Neurons. The Journal of neuroscience: the official journal of the Society for Neuroscience 37, 8678–8687. 10.1523/JNEUROSCI.0798-17.2017 [doi].

19. Gropp, E., Shanabrough, M., Borok, E., Xu, A.W., Janoschek, R., Buch, T., Plum, L., Balthasar, N., Hampel, B., Waisman, A., et al. (2005). Agouti-related peptide-expressing neurons are mandatory for feeding. Nature neuroscience 8, 1289–1291. nn1548 [pii].

20. Betley, J.N., Xu, S., Cao, Z.F.H., Gong, R., Magnus, C.J., Yu, Y., and Sternson, S.M. (2015). Neurons for hunger and thirst transmit a negative-valence teaching signal. Nature 521, 180–185. 10.1038/nature14416 [doi].

21. Bai, L., Mesgarzadeh, S., Ramesh, K.S., Huey, E.L., Liu, Y., Gray, L.A., Aitken, T.J., Chen, Y., Beutler, L.R., Ahn, J.S., et al. (2019). Genetic Identification of Vagal Sensory Neurons That Control Feeding. Cell 179, 1129–1143.e1123. S0092-8674(19)31181-X [pii].

22. Rossi, M.A., Basiri, M.L., McHenry, J.A., Kosyk, O., Otis, J.M., van den Munkhof, H.E., Bryois, J., Hubel, C., Breen, G., Guo, W., et al. (2019). Obesity remodels activity and transcriptional state of a lateral hypothalamic brake on feeding. Science (New York, N.Y.) 364, 1271–1274. 10.1126/science.aax1184 [doi].

23. Sternson, S.M., and Eiselt, A.-K. (2017). Three Pillars for the Neural Control of Appetite. Annual Review of Physiology 79, 401–423. 10.1146/annurev-physiol-021115-104948.

24. Goldstein, N., McKnight, A.D., Carty, J.R.E., Arnold, M., Betley, J.N., and Alhadeff, A.L. (2021). Hypothalamic detection of macronutrients via multiple gut-brain pathways. Cell metabolism. 10.1016/j.cmet.2020.12.018. S1550-4131(20)30716-6 [pii].

25. Korgan, A.C., Oliveira-Abreu, K., Wei, W., Martin, S.L.A., Bridges, Z.J.D., Leal-Cardoso, J.H., Kaczorowski, C.C., and O’Connell, K.M.S. (2023). High sucrose consumption decouples intrinsic and synaptic excitability of AgRP neurons without altering body weight. International Journal of Obesity 47, 224–235. 10.1038/s41366-023-01265-w.

26. Mitchell, C.S., Goodman, E.K., Tedesco, C.R., Nguyen, K., Zhang, L., Herzog, H., and Begg, D.P. (2022). The Effect of Dietary Fat and Sucrose on Cognitive Functioning in Mice Lacking Insulin Signaling in Neuropeptide Y Neurons. Frontiers in Physiology 13. 10.3389/fphys.2022.841935.

27. Chen, G.-C., Huang, C.-Y., Chang, M.-Y., Chen, C.-H., Chen, S.-W., Huang, C.-j., and Chao, P.-M. (2011). Two unhealthy dietary habits featuring a high fat content and a sucrose-containing beverage intake, alone or in combination, on inducing metabolic syndrome in Wistar rats and C57BL/6J mice. Metabolism 60, 155–164. https://doi.org/10.1016/j.metabol.2009.12.002.

28. Sakamoto, E., Seino, Y., Fukami, A., Mizutani, N., Tsunekawa, S., Ishikawa, K., Ogata, H., Uenishi, E., Kamiya, H., Hamada, Y., et al. (2012). Ingestion of a moderate high-sucrose diet results in glucose intolerance with reduced liver glucokinase activity and impaired glucagon-like peptide-1 secretion. Journal of Diabetes Investigation 3, 432–440. https://doi.org/10.1111/j.2040-1124.2012.00208.x.

29. Leibel, R.L., Rosenbaum, M., and Hirsch, J. (1995). Changes in energy expenditure resulting from altered body weight. The New England journal of medicine 332, 621–628. 10.1056/NEJM199503093321001 [doi].

30. Fothergill, E., Guo, J., Howard, L., Kerns, J.C., Knuth, N.D., Brychta, R., Chen, K.Y., Skarulis, M.C., Walter, M., Walter, P.J., and Hall, K.D. (2016). Persistent metabolic adaptation 6 years after “The Biggest Loser” competition. Obesity (Silver Spring, Md.) 24, 1612–1619. 10.1002/oby.21538 [doi].

31. Myers, M.G., Jr., Leibel, R.L., Seeley, R.J., and Schwartz, M.W. (2010). Obesity and leptin resistance: distinguishing cause from effect. Trends in endocrinology and metabolism: TEM 21, 643–651. 10.1016/j.tem.2010.08.002 [doi].

32. Hwang, E., Scarlett, J.M., Baquero, A.F., Bennett, C.M., Dong, Y., Chau, D., Brown, J.M., Mercer, A.J., Meek, T.H., Grove, K.L., et al. (2022). Sustained inhibition of NPY/AgRP neuronal activity by FGF1. JCI Insight 7. 10.1172/jci.insight.160891.

33. Briggs, D.I., Lockie, S.H., Wu, Q., Lemus, M.B., Stark, R., and Andrews, Z.B. (2013). Calorie-restricted weight loss reverses high-fat diet-induced ghrelin resistance, which contributes to rebound weight gain in a ghrelin-dependent manner. Endocrinology 154, 709–717. 10.1210/en.2012-1421 [doi].

34. Schmitz, J., Evers, N., Awazawa, M., Nicholls, H.T., Bronneke, H.S., Dietrich, A., Mauer, J., Bluher, M., and Bruning, J.C. (2016). Obesogenic memory can confer long-term increases in adipose tissue but not liver inflammation and insulin resistance after weight loss. Molecular metabolism 5, 328–339. S2212-8778(15)00231-8 [pii].

35. Molly, M., Alan de, A., Macarena, V., Mingxin, Y., Arashdeep, S., Isadora, B., Nikhil, U., Brandon, W., and Guillaume de, L. (2022). Labeled lines for fat and sugar reward combine to promote overeating. bioRxiv, 2022.2008.2009.503218. 10.1101/2022.08.09.503218.

36. Edwin Thanarajah, S., DiFeliceantonio, A.G., Albus, K., Kuzmanovic, B., Rigoux, L., Iglesias, S., Hanßen, R., Schlamann, M., Cornely, O.A., Brüning, J.C., et al. (2023). Habitual daily intake of a sweet and fatty snack modulates reward processing in humans. Cell Metabolism 35, 571–584.e576. https://doi.org/10.1016/j.cmet.2023.02.015.

37. DiFeliceantonio, A.G., Coppin, G., Rigoux, L., Edwin Thanarajah, S., Dagher, A., Tittgemeyer, M., and Small, D.M. (2018). Supra-Additive Effects of Combining Fat and Carbohydrate on Food Reward. Cell Metabolism 28, 33–44.e33. https://doi.org/10.1016/j.cmet.2018.05.018.

38. Briggs, D.I., Lemus, M.B., Kua, E., and Andrews, Z.B. (2011). Diet-induced obesity attenuates fasting-induced hyperphagia. Journal of neuroendocrinology 23, 620–626. 10.1111/j.1365-2826.2011.02148.x [doi].

39. Garfield, A.S., Shah, B.P., Burgess, C.R., Li, M.M., Li, C., Steger, J.S., Madara, J.C., Campbell, J.N., Kroeger, D., Scammell, T.E., et al. (2016). Dynamic GABAergic afferent modulation of AgRP neurons. Nature neuroscience 19, 1628–1635. 10.1038/nn.4392 [doi].

40. Smith, M.A., Choudhury, A.I., Glegola, J.A., Viskaitis, P., Irvine, E.E., de Campos Silva, P.C.C., Khadayate, S., Zeilhofer, H.U., and Withers, D.J. (2020). Extrahypothalamic GABAergic nociceptin–expressing neurons regulate AgRP neuron activity to control feeding behavior. The Journal of Clinical Investigation 130, 126–142. 10.1172/JCI130340.

41. Berrios, J., Li, C., Madara, J.C., Garfield, A.S., Steger, J.S., Krashes, M.J., and Lowell, B.B. (2021). Food cue regulation of AGRP hunger neurons guides learning. Nature 595, 695–700. 10.1038/s41586-021-03729-3.

42. Grzelka, K., Wilhelms, H., Dodt, S., Dreisow, M.-L., Madara, J.C., Walker, S.J., Wu, C., Wang, D., Lowell, B.B., and Fenselau, H. (2023). A synaptic amplifier of hunger for regaining body weight in the hypothalamus. Cell Metabolism 35, 770–785.e775. https://doi.org/10.1016/j.cmet.2023.03.002.

43. Briggs, D.I., Enriori, P.J., Lemus, M.B., Cowley, M.A., and Andrews, Z.B. (2010). Diet-induced obesity causes ghrelin resistance in arcuate NPY/AgRP neurons. Endocrinology 151, 4745–4755. 10.1210/en.2010-0556 [doi].

44. Zigman, J.M., Bouret, S.G., and Andrews, Z.B. (2016). Obesity Impairs the Action of the Neuroendocrine Ghrelin System. Trends in endocrinology and metabolism: TEM 27, 54–63. S1043-2760(15)00193-9 [pii].

45. Qi, Y., Lee, N.J., Ip, C.K., Enriquez, R., Tasan, R., Zhang, L., and Herzog, H. (2023). Agrp-negative arcuate NPY neurons drive feeding under positive energy balance via altering leptin responsiveness in POMC neurons. Cell Metabolism 35, 979–995.e977. https://doi.org/10.1016/j.cmet.2023.04.020.

46. Ahn, B.H., Kim, M., and Kim, S.-Y. (2022). Brain circuits for promoting homeostatic and non-homeostatic appetites. Experimental & Molecular Medicine 54, 349–357. 10.1038/s12276-022-00758-4.

47. González-Padilla, E., A. Dias, J., Ramne, S., Olsson, K., Nälsén, C., and Sonestedt, E. (2020). Association between added sugar intake and micronutrient dilution: a cross-sectional study in two adult Swedish populations. Nutrition & Metabolism 17, 15. 10.1186/s12986-020-0428-6.

48. Grech, A., Sui, Z., Rangan, A., Simpson, S.J., Coogan, S.C.P., and Raubenheimer, D. (2022). Macronutrient (im)balance drives energy intake in an obesogenic food environment: An ecological analysis. Obesity 30, 2156–2166. https://doi.org/10.1002/oby.23578.

49. Ueno, A., Lazaro, R., Wang, P.Y., Higashiyama, R., Machida, K., and Tsukamoto, H. (2012). Mouse intragastric infusion (iG) model. Nature protocols 7, 771–781. 10.1038/nprot.2012.014 [doi].

50. Chen, Y., Lin, Y.C., Zimmerman, C.A., Essner, R.A., and Knight, Z.A. (2016). Hunger neurons drive feeding through a sustained, positive reinforcement signal. eLife 5, 10.7554/eLife.18640. [doi].

51. Liang, W., Menke, A.L., Driessen, A., Koek, G.H., Lindeman, J.H., Stoop, R., Havekes, L.M., Kleemann, R., and van den Hoek, A.M. (2014). Establishment of a general NAFLD scoring system for rodent models and comparison to human liver pathology. PLoS One 9, e115922. 10.1371/journal.pone.0115922.

